# Achieving dendritic cell subset-specific targeting *in vivo* by site-directed conjugation of targeting antibodies to nanocarriers

**DOI:** 10.1101/2021.07.14.452311

**Authors:** Johanna Simon, Michael Fichter, Gabor Kuhn, Maximilian Brückner, Cinja Kappel, Jenny Schunke, Stephan Grabbe, Katharina Landfester, Volker Mailänder

## Abstract

The major challenge of nanocarrier-based anti-cancer vaccination approaches is the targeted delivery of antigens and immunostimulatory agents to cells of interest, such as specific subtypes of dendritic cells (DCs), in order to induce robust antigen-specific anti-tumor responses. An undirected cell and body distribution of nanocarriers can lead to unwanted delivery to other immune cell types like macrophages reducing the vaccine efficacy. An often-used approach to overcome this issue is the surface functionalization of nanocarriers with targeting moieties, such as antibodies, mediating cell type-specific interaction. Numerous studies could successfully prove the targeting efficiency of antibody-conjugated carrier systems *in vitro*, however, most of them failed when targeting DCs *in vivo* that is partly due to cells of the reticuloendothelial system unspecifically clearing nanocarriers from the blood stream via Fc receptor ligation.

Therefore, this study shows a surface functionalization strategy to site-specifically attach antibodies in an orientated direction onto the nanocarrier surface. Different DC-targeting antibodies, such as anti-CD11c, anti-CLEC9A, anti-DEC205 and anti-XCR1, were conjugated to the nanocarrier surface at their Fc domains. Anti-mouse CD11c antibody-conjugated nanocarriers specifically accumulated in the targeted organ (spleen) over time. Additionally, antibodies against CD11c and CLEC9A proved to specifically direct nanocarriers to the targeted DC subtype, conventional DCs type 1.

In conclusion, site-directed antibody conjugation to nanocarriers is essential in order to avoid unspecific uptake by non-target cells while achieving antibody-specific targeting of DC subsets. This novel conjugation technique paves the way for the development of antibody-functionalized nanocarriers for DC-based vaccination approaches in the field of cancer immunotherapy.

## Introduction

Nanocarriers (NC) applied as drug delivery vehicles protect the cargo in an unprecedented way, as exemplified these days with the newly approved preventive SARS-CoV-2 vaccines developed by BioNtech or Moderna loaded with highly fragile loads like mRNA. This revolutionizes the current strategy for preventing viral diseases or treating cancer [1-5]. Also an unwanted body distribution can be skewed towards a more desired one. To this end, a specific transport of the drug to cells of interest via the nanocarrier can be achieved through the functionalization of the nanocarrier surface with targeting ligands such as antibodies specific for surface antigens [6, 7]. Antibody-conjugated nanocarriers have the potential to improve the drug selectivity, reduce systemic side effects, and increase the therapeutic potential [8].

Already in the 90s, the first nanocarrier system (DOXIL®), a polyethylene glycol (PEG)- functionalized liposomal formulation of doxorubicin, was approved by the FDA for treating cancer [9, 10]. To date, however, no antibody-conjugated nanocarrier system has been clinically approved. Even though numerous reports could successfully prove the targeting specificity of antibody-conjugated nanocarriers *in vitro* [11, 12], the majority of these studies failed *in vivo* (see e.g. [13, 14]). A major factor responsible for these discrepancies is the phenomenon of blood protein adsorption on the nanocarrier surface following intravenous injection forming the biomolecular corona [15, 16]. Due to this process, the targeting ligand can be completely covered up by corona proteins preventing an interaction between the antibody conjugated onto the nanocarrier and its targeted cell receptor [17, 18]. A second confounding factor which enhances unspecific uptake by especially macrophages is binding of the Fc part of antibodies to Fc receptors. We have shown before that this can be avoided by a site-directed coupling strategy [19].

To overcome current challenges, this study presents a surface functionalization strategy for the attachment of dendritic cell-specific antibodies onto a nanocarrier surface allowing a targeted cell interaction *in vivo*. The effect of protein corona formation was studied both *in vitro* as well as *in vivo*, and the targeting properties of the designed antibody-conjugated nanocarriers were not affected. This approach can be transferred to a broad range of nanocarriers and antibodies. As a proof of concept, anti-mouse CD11c antibody was conjugated to the surface of magnetic nanoparticles. The CD11c receptor is expressed by all subtypes of dendritic cells [20] present in the liver [21] and the spleen [22]. Targeting dendritic cells is of great interest in the field of cancer immunotherapy [23], as DCs play a crucial role in activating the immune system to fight cancer [24] and other diseases [25]. CLEC9A and DEC205 define subgroups of CD11c+ DCs. But not all DCs support an anti-viral or anti-tumor response in the same effective way. CLEC9A+ and DEC205+ DCs are thought to define subgroups which are crucial for an effective immune response. This study demonstrates that anti-mouse CD11c-, CLEC9A-, as well as DEC205-functionalized nanocarriers specifically bind to distinct subtypes of dendritic cells *in vivo*, whereas nanocarriers conjugated to an IgG control antibody exhibited no interaction with the targeted cell population.

## Results and discussion

To maintain the targeting properties of the antibody and hereby enable an effective transport of the nanocarrier *in vivo*, a site-specific antibody functionalization strategy was chosen [19]. Therefore, the antibody is enzymatically modified in a two-step reaction at the Fc part with a maximum of four azide groups (two N_3_-groups per heavy chain). This functionalization strategy takes advantage of the characteristic N-glycosylation pattern at the Fc region (CH_2_ domain at the amino acid asparagine 297 (N297)), which is present at all immunoglobulins. First, galactose residues were enzymatically removed and, in a second step, using a transferase the azide-modified sugar component N-azidoacetylglucosamine (GalNAz) was attached resulting in a site-specific modification (Figure 1A). With this strategy the antigen binding region of the antibody remains unmodified, therefore, mediating specific binding towards the targeted cell receptor.

**Figure 1.**
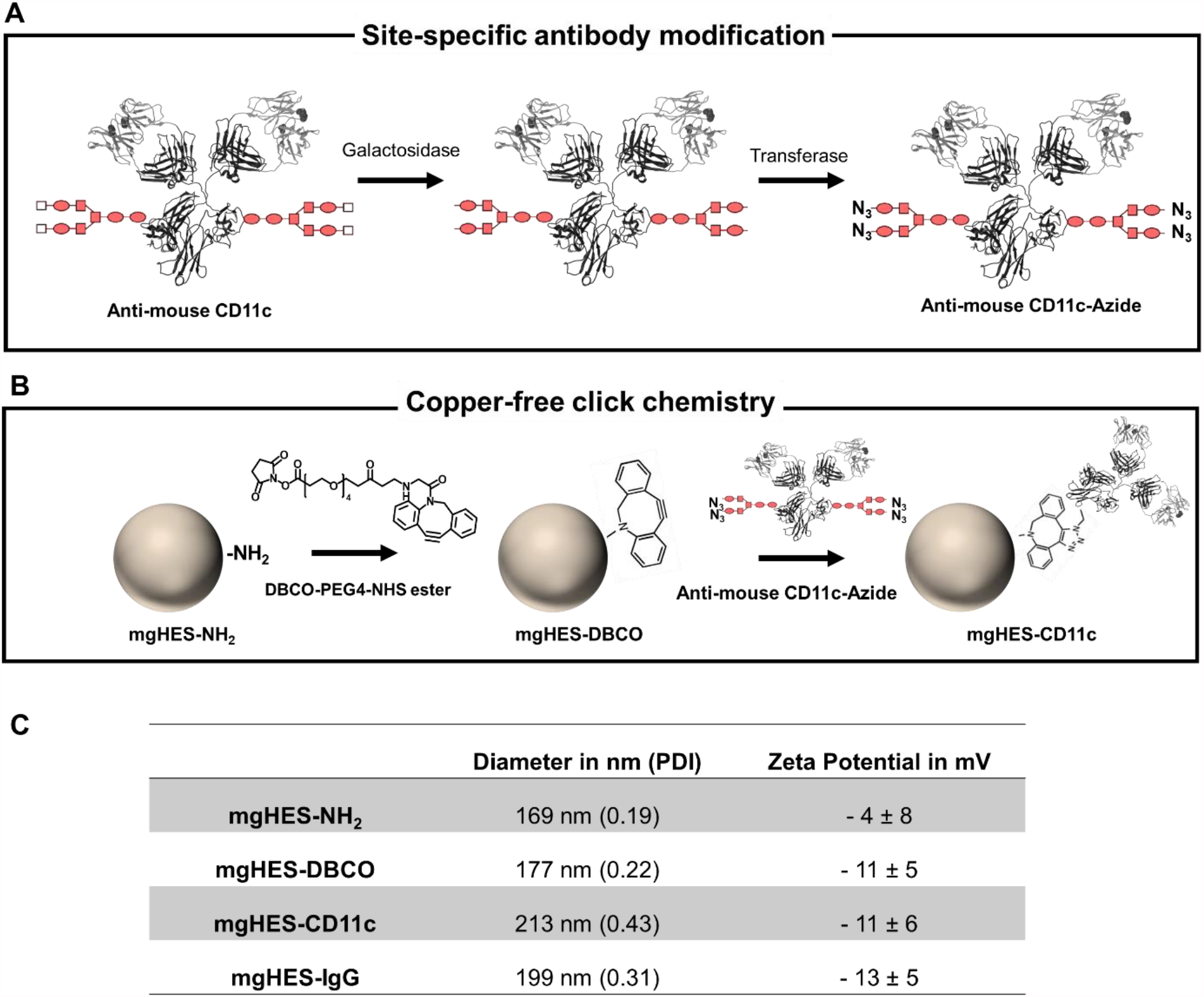
Site-specific antibody modification and the functionalization of nanocarriers. **A)** Antibodies were enzymatically modified with a maximum of four azides on the Fc region. **B)** Nanocarriers were functionalized with a DBCO linker via NHS-chemistry and further via copper-free click reaction the antibody was attached on the nanocarrier surface. **C)** Physico-chemical properties of all nanocarriers before and after functionalization.

The mgHES nanocarriers carrying amino-groups on the surface were functionalized via NHS-chemistry with an alkyne-modified DBCO-PEG-linker (Figure 1 B). Finally, the azide-modified antibody is coupled via the copper-free click reaction to alkyne-functionalized DBCO-functionalized nanocarriers representing quantitative [26] and biorthogonal conjugation process [27, 28] (Figure 1 B). Altogether, this functionalization strategy allows the precise attachment of an antibody to the surface of a nanocarrier. In addition, this attachment protocol can be transferred to any nanocarrier with amino-groups on the surface.

Non-functionalized and functionalized nanocarriers were physico-chemically characterized using dynamic light scattering and zeta potential measurements. Antibody coupling resulted in a minor size increase (∼30 nm) and slightly reduced zeta potential (Figure 1 C). For the initial characterization of antibody functionalization, an anti-mouse CD11c antibody binding to CD11c-expressing dendritic cells was used. Dendritic cells represent key players for the induction of antigen-specific anti-tumor responses in vaccination approaches and are of great interest as a target cell population using nanocarrier-based antigen delivery [29]. Additionally, an isotype control antibody was selected to demonstrate the cellular targeting specificity of the anti-mouse CD11c-modified nanocarriers.

Wild type C57BL/6 mice were treated with antibody-modified nanocarriers (1 mg) or with PBS as a control. The blood was isolated after 1 min, 10 min, 60 min, or 120 min to determine the blood circulation time of the antibody-modified nanocarriers (Figure 2 B). Next to this, the nanocarriers remaining in the blood were magnetically recovered to determine the proteins, attached onto the nanocarrier surface after *in vivo* circulation (Figure 2 A). At distinct time points, mice were sacrificed, organs were isolated and analyzed using the IVIS® SpectrumCT imager to investigate the biodistribution.

**Figure 2.**
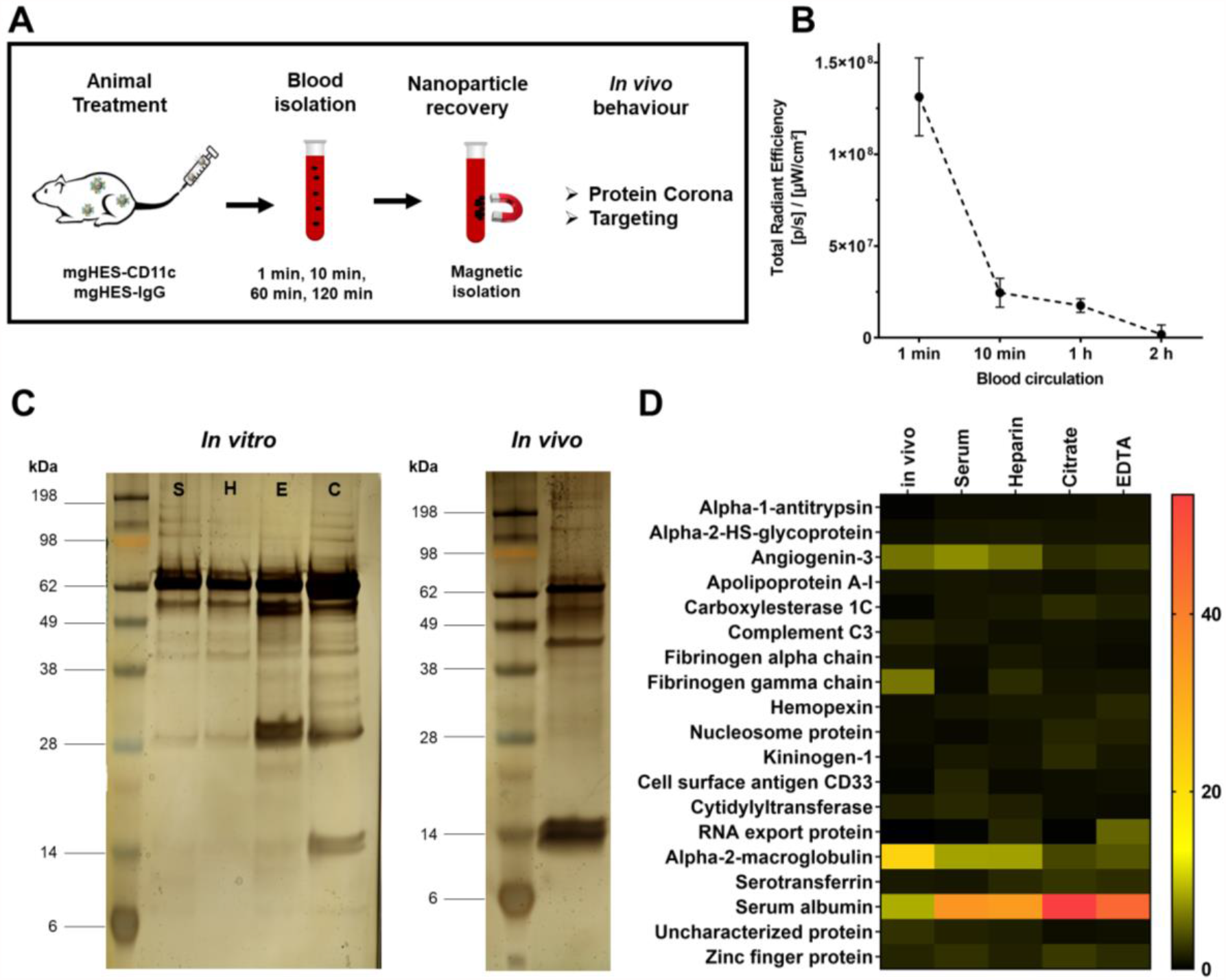
*In vivo* protein corona formation. **A)** Experimental workflow **B)** Blood circulation time of antibody-functionalized nanocarriers. Mice were treated with mgHES-CD11c nanocarriers or PBS as a control. At different time points blood was isolated and analyzed via the IVIS® SpectrumCT. Background fluorescence measured from PBS-treated mice was subtracted. **C)** SDS-PAGE of the protein corona of antibody-functionalized nanocarriers incubated under *in vitro* conditions for 1 min (S = Serum, H = Heparin Plasma, E = EDTA Plasma, C = Citrate Plasma) or after recovery from the blood stream after 1 min (*in vivo*). **D)** LC-MS analysis of the protein corona of antibody-functionalized nanocarriers. The relative amount of each protein in % is calculated based on the total amount of all identified proteins determined in fmol. *n* = 3, for all *in vivo* samples.

It has been shown that the adsorption of corona proteins onto the surface of antibody-modified nanocarriers can significantly affect their binding properties to the targeted cells *in* vitro [30, 31]. However, only few studies have characterized the protein corona formation of nanocarriers after *in vivo* application and it has been shown that there is a significant difference in the protein composition compared to the *in vitro* situation [32-34]. Therefore, the proteins adsorbing to the nanocarriers in the blood stream *in vivo* were analyzed via SDS-PAGE and proteomic high-resolution mass spectrometry. Additionally, the protein pattern after *in vivo* circulation was compared to the protein corona pattern after *in vitro* incubation with serum or plasma. Most importantly, the protein analyses revealed a significant difference in the corona proteome of nanocarriers incubated under *in vitro* conditions compared to the *in vivo* situation (Figure 2 C and D). This effect was observed for both control (mgHES-IgG) and targeting nanocarriers (mgHES-CD11c) under either *in vitro* (Figure S1) or *in vivo* (Figure S2) conditions. For all nanocarriers, alpha-2 macroglobulin was enriched in the protein corona after *in vivo* circulation, whereas primarily serum albumin was bound to nanocarriers after *in vitro* incubation (Figure S2). These results indicate that studies characterizing the protein corona forming after in vitro incubation do not fully reflect the complex situation present in biological fluids such as the blood stream or interstitial fluid. Additionally, these findings give hints to explain the great discrepancy between successful *in vitro* studies and failing *in vivo* experiments aiming to target specific cell populations. To overcome this issue, we developed a method to mimic the *in vivo* corona in a more precise fashion [32]. Therefore, nanocarriers were incubated with mouse whole blood and the protein corona was prepared (indicated as *ex vivo*). High-resolution mass spectrometry demonstrated that the protein corona pattern after *ex vivo* incubation is highly comparable to the *in vivo* pattern (Figure S2). This finding indicates that the *ex vivo* strategy allows to mimic the *in vivo* protein corona. A full list of all identified proteins under *in vitro, ex vivo*, and *in vivo* conditions for both nanocarriers is provided in a separate Excel File.

A five-step protocol was implemented investigating the efficiency of antibody-functionalized nanocarriers to specifically address target cells *in vivo*. In a first step, nanocarriers recovered from treated mice after *in vivo* blood circulation were incubated with dendritic cells (DC2.4) expressing the CD11c receptor on their cell surface *in vitro* (Figure 3 A). These experiments served to delineate the effects of the *in vivo* protein corona upon the antibody-functionalized nanocarriers’ targeting properties *in vitro*. As a control, nanocarriers were left untreated (w/o corona) or pre-treated with either serum or plasma to allow *in vitro* protein corona formation or with whole blood for *ex vivo* incubation (w/ corona). The cellular binding affinity of all nanocarriers was analyzed. Flow cytometry revealed that all anti-CD11c-functionalized nanocarriers specifically bound to DC2.4 cells irrelevant of pre-treatment, whereas the IgG control nanocarriers showed a significantly reduced cell binding (Figure 3 A). Comparable results were also obtained for anti-CD11c-functionalized nanocarriers after *ex vivo* blood incubation.

**Figure 3.**
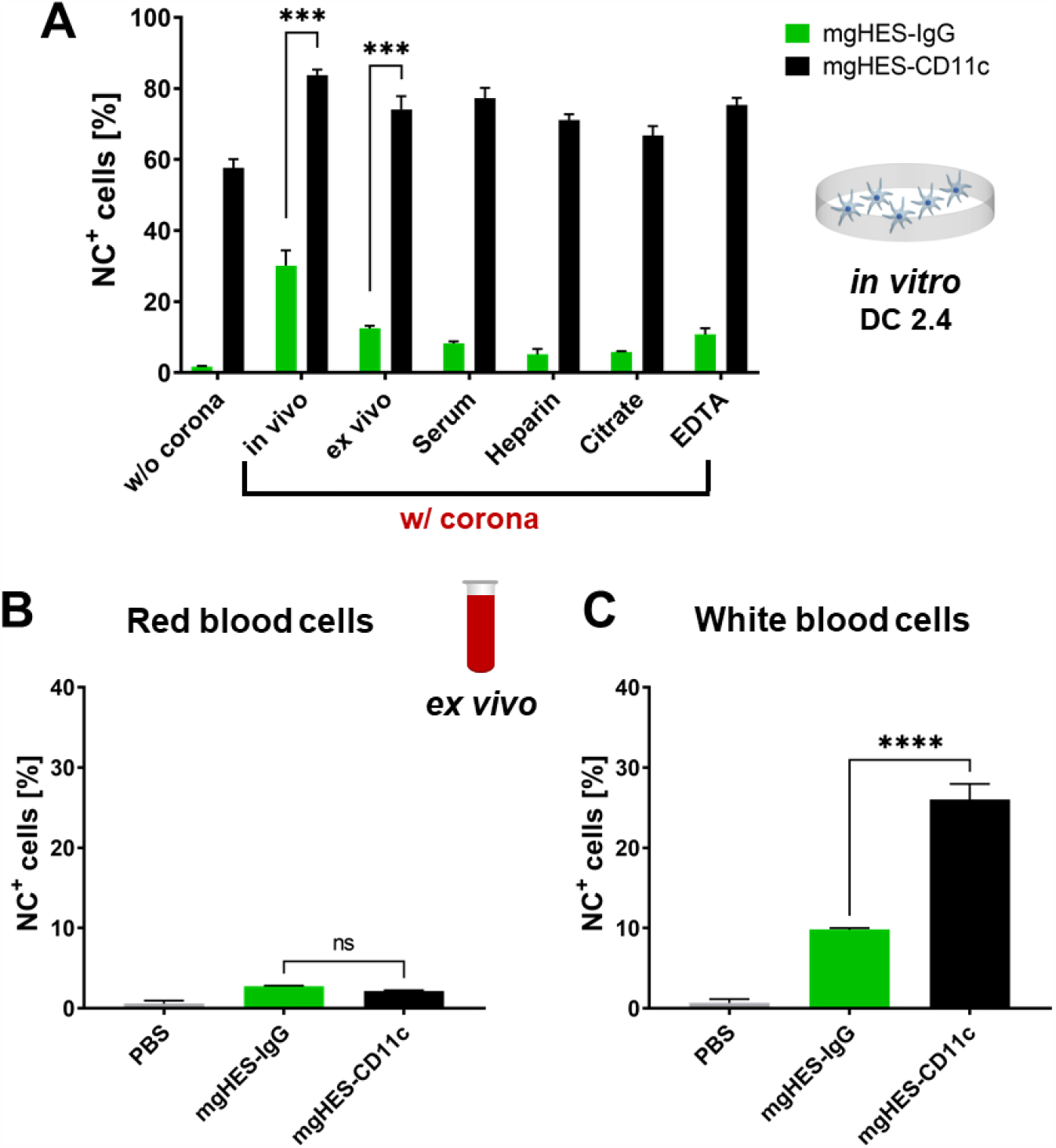
*In vitro* and *ex vivo* targeting properties of anti-CD11c-functionalized nanocarriers. **A)** DC2.4 cells were incubated with antibody-functionalized nanocarriers (37.5 µg mL^-1^) without protein corona coating (indicated as w/o corona), after *in vivo* circulation (1 min, indicated as *in vivo* w/ corona), after *ex vivo* protein coating for 1 min (indicated as *ex vivo*), or after *in vitro* protein coating for 2 h (indicated as Serum, Heparin, Citrate, EDTA w/ corona). Data represent mean ± SD. *In vivo* and *ex vivo* protein corona-coated mgHES-CD11c were compared to mgHES-IgG, respectively, and significance was given with *p* < 0.05 (Welch’s *t* test). *p* < 0.001***. **B+C)** Interaction of whole blood components with antibody functionalized nanocarriers. The whole blood was analyzed via flow cytometry and the interaction of antibody-functionalized nanocarriers with red blood cells (**B**) and white blood cells (**C**) was measured using flow cytometry. Data represent mean ± SD and significance was given with *p* < 0.05 (one-way ANOVA). *ns* = not significant, *p* < 0.0001****.

This finding proves that even after *in vivo* circulation or *ex vivo* blood incubation the antibodies on the nanocarrier surface retained their targeting properties. It also highlights that even in the presence of the protein corona the antibody-functionalized nanocarriers are able to bind to the targeted cells.

In a second step, an *ex vivo* approach was developed. Therefore, mouse whole blood was incubated with antibody-functionalized nanocarriers. White and red blood cells were differentiated in whole blood by flow cytometry due to distinct light-scattering properties [35]. A detailed protocol is provided in the method section. Interactions of anti-CD11c-functionalized nanocarriers with erythrocytes could not be observed (Figure 3 B). This is of great importance as it has been shown that some nanocarriers favor the interaction with red blood cells and hereby induce hemolysis and severe side effects [36, 37].

Furthermore, anti-CD11c-functionalized nanocarriers specifically bound to white blood cells composed of a mixture of different cell types including CD11c^+^ cells, while the frequency of mgHES-IgG^+^ cells was significantly reduced [38].

In a third step, the targeting efficacy of the anti-CD11c-functionalized nanocarriers was investigated on organ level. Therefore, mgHES-CD11c- or mgHES-IgG-treated mice were sacrificed and all relevant organs were dissected. Liver, spleen, lung and kidneys were subsequently analyzed for nanocarrier accumulation using small animal fluorescence imaging (Figure 4).

**Figure 4.**
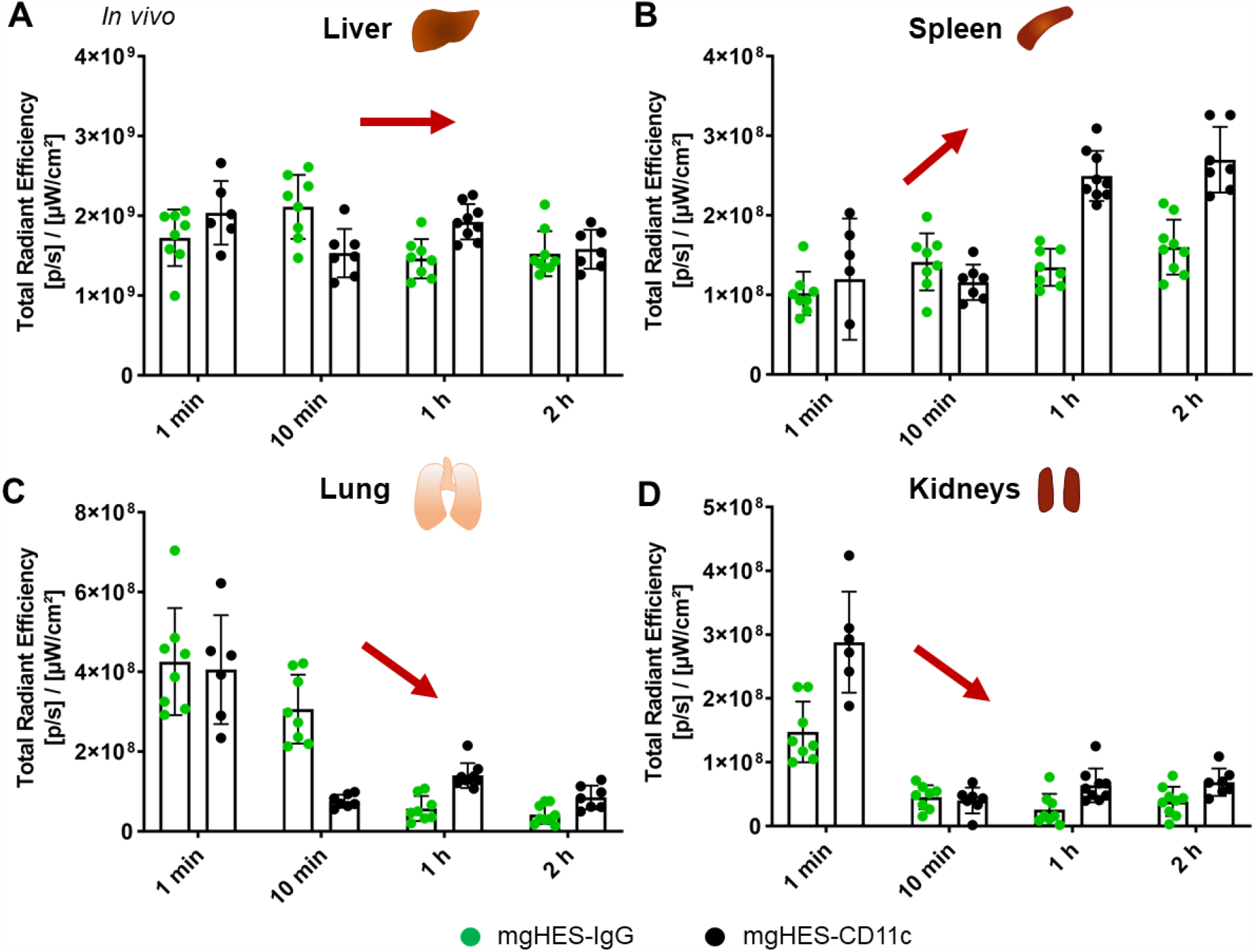
Biodistribution of antibody-functionalized mgHES nanocarriers into different organs over time. **A) – D)** Animals were intravenously injected with mgHES-IgG or mgHES-CD11c nanocarriers (1 mg). Liver (A), spleen (B), lung (C), and kidneys (D) were dissected at different time points post injection (1 min – 2 h) and imaged with IVIS® SpectrumCT. The fluorescence intensity of all organs was analyzed and background fluorescence of each respective organ was subtracted based on mice injected with PBS as a control.

Foremost, both nanocarrier formulations primarily accumulated in the liver (Figure 4 A). This phenomenon has already been reported for a great number of studies [38-41] and is based on the physiological function of the liver as a filter organ clearing the bloodstream from foreign particulate material [42]. This can lead to off-target effects and unwanted side effects [42, 43]. Nevertheless, CD11c^+^ dendritic cells also reside within the liver. Therefore, the binding of anti-CD11c-functionalized nanocarriers towards CD11c^+^ and CD11c^-^ non-parenchymal liver cells was determined in order to investigate the specificity of nanocarrier targeting within the liver. (Figure 5).

**Figure 3.**
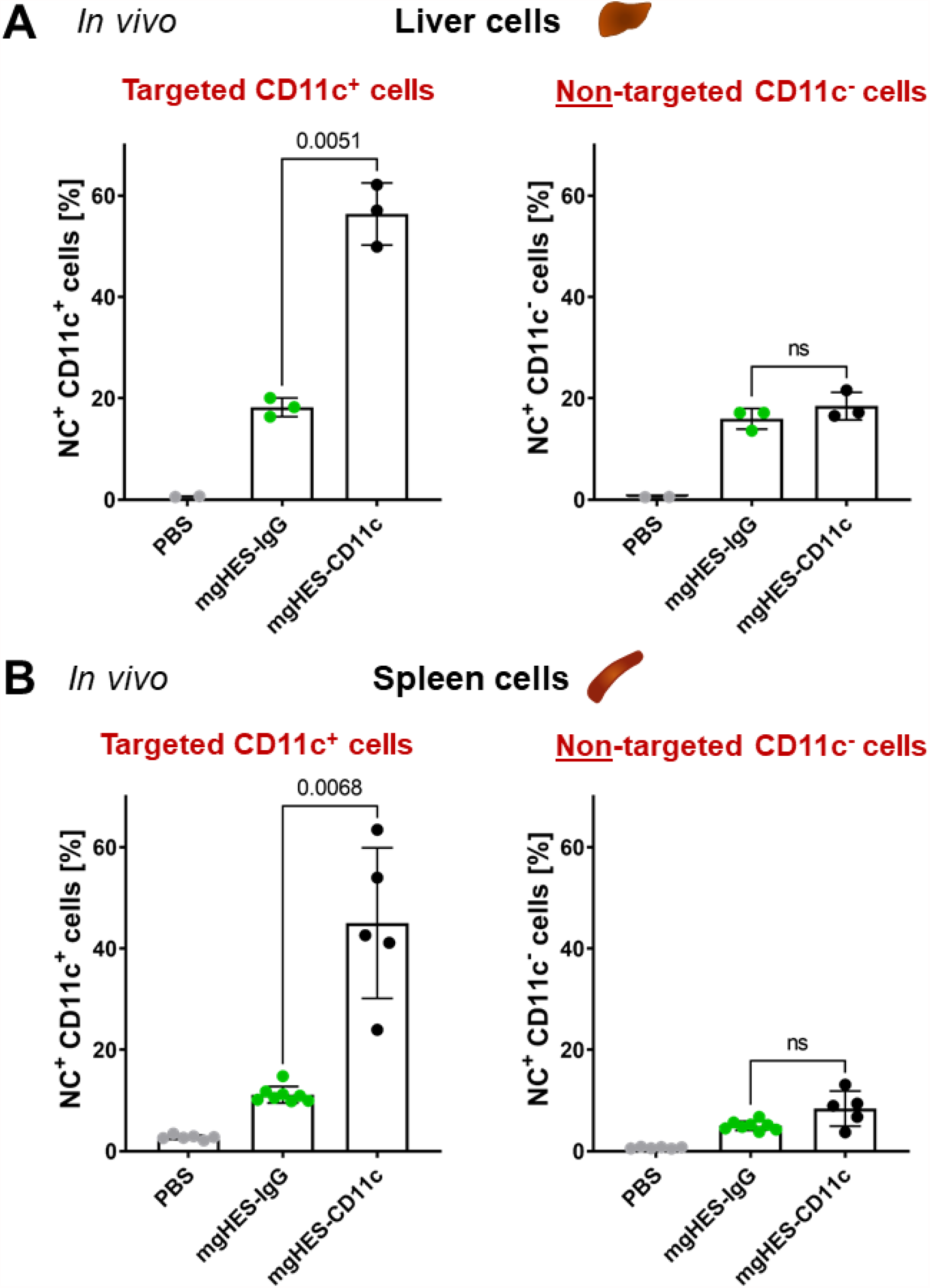
Targeting CD11c^+^ liver and spleen cells *in vivo*. Antibody-functionalized nanocarriers were intravenously injected into mice. Non-parenchymal liver cells **(A)** were isolated 2 h and splenocytes **(B)** 1 min after injection and nanocarrier uptake by CD11c^+^ or CD11c^-^ cells was analyzed using flow cytometry. Data represent mean ± SD. mgHES-CD11c nanocarriers were compared to the IgG control (mgHES-IgG) and significance was given with *p* < 0.05 using a Welch’s *t* test. *ns* = not significant. Individual *p* values are indicated in the graph.

For the spleen, additionally harboring CD11c^+^ cells, the fluorescence signal of the anti-CD11c-functionalized nanocarriers steadily increased from 1 min to 120 min. following injection, while no significant increase in IgG control nanocarriers could be observed (Figure 4 B). This finding indicates an anti-CD11c-specific accumulation of nanocarriers in the targeted organ over time. In contrast, the nanocarrier-based fluorescence signal in the non-targeted organs (lung and kidney) decreased over time (Figure 4 C and D).

As described above, in a fourth step (Figure 5), the targeting efficiency of the antibody-functionalized nanocarriers *in vivo* towards CD11c^+^ and CD11^-^ liver or spleen cells was investigated. This allowed determining the binding specificity of anti-CD11c-functionalized nanocarriers and indicated binding to targeted or non-targeted cells. Flow cytometric analyses revealed that anti-CD11c-functionalized nanocarriers specifically bound to CD11c^+^ cells, both in the liver and in the spleen, while the frequency of cells positive for IgG control nanocarriers was significantly reduced in both organs (Figure 5 A and B and Figure S3). Targeting specificity *in vivo* was additionally proved by analyzing CD11c^-^ cells showing no significant differences in binding of either anti-CD11c- or IgG control nanocarriers (Figure 5 A and B and Figure S3). In conclusion, this proves that the here presented anti-CD11c-functionalized nanocarriers were able to reach the targeted cells *in vivo*.

In the final step, additional DC-specific antibodies were conjugated onto the mgHES nanocarriers in order to evaluate their capacity for nanocarrier-based targeting of relevant DC subpopulations. CD8α^+^ conventional DCs type 1 (cDC1) are of great interest for the targeted delivery of antigens in vaccination approaches due to their high cross-presenting potential and efficiency to prime anti-tumor CD8^+^ T cells [29, 44]. Next to the pan DC marker CD11c, cDC1 cells express XCR1, CLEC9A, and DEC205 on their cell surface, which have been exploited as targeting receptors in vaccination studies either as antibody-antigen fusion proteins or as antibody-functionalized nanoparticles [45-49]. Therefore, antibodies against these receptors present on cDC1 cells were conjugated onto mgHES nanocarriers using the method described above and their targeting efficacy towards different cell types within the spleen was analyzed. IgG control and anti-CD11c-functionalized nanocarriers were applied as negative and positive controls, respectively, and led to similar frequencies of NC^+^ CD11c^+^ (Figure 6 A) and CD11c^-^ (Figure 6 B) cells compared to the results described above (Figure 5 B). In addition to the analysis of pan DC populations (CD11c^+^; Figure 6 A), anti-CD11c-functionalized nanocarriers bound to all analyzed CD11c^+^ DC subtypes in an elevated manner (Figure 6 C - E). Notably, functionalization of nanocarriers with anti-XCR1 did not result in an elevated uptake by any of the cell types investigated (Figure 6). While anti-DEC205 nanocarriers did not improve binding to cDC1, plasmacytoid DCs and cDC2 showed an increased binding (Figure 6 D and E). DEC205 is often exploited as a targeting receptor for cDC1, however, there is evidence that also cDC2 express this endocytic C-type lectin molecule [50, 51]. The most pronounced elevation in binding of cDC1 and plasmacytoid DCs was achieved with anti-CLEC9A-functionalized nanocarriers that even exceeded the effects induced with CD11c targeting (Figure 6 C and E). CLEC9A has been found to be differentially expressed on conventional DCs type 1, but also on plasmacytoid [52].

**Figure 6.**
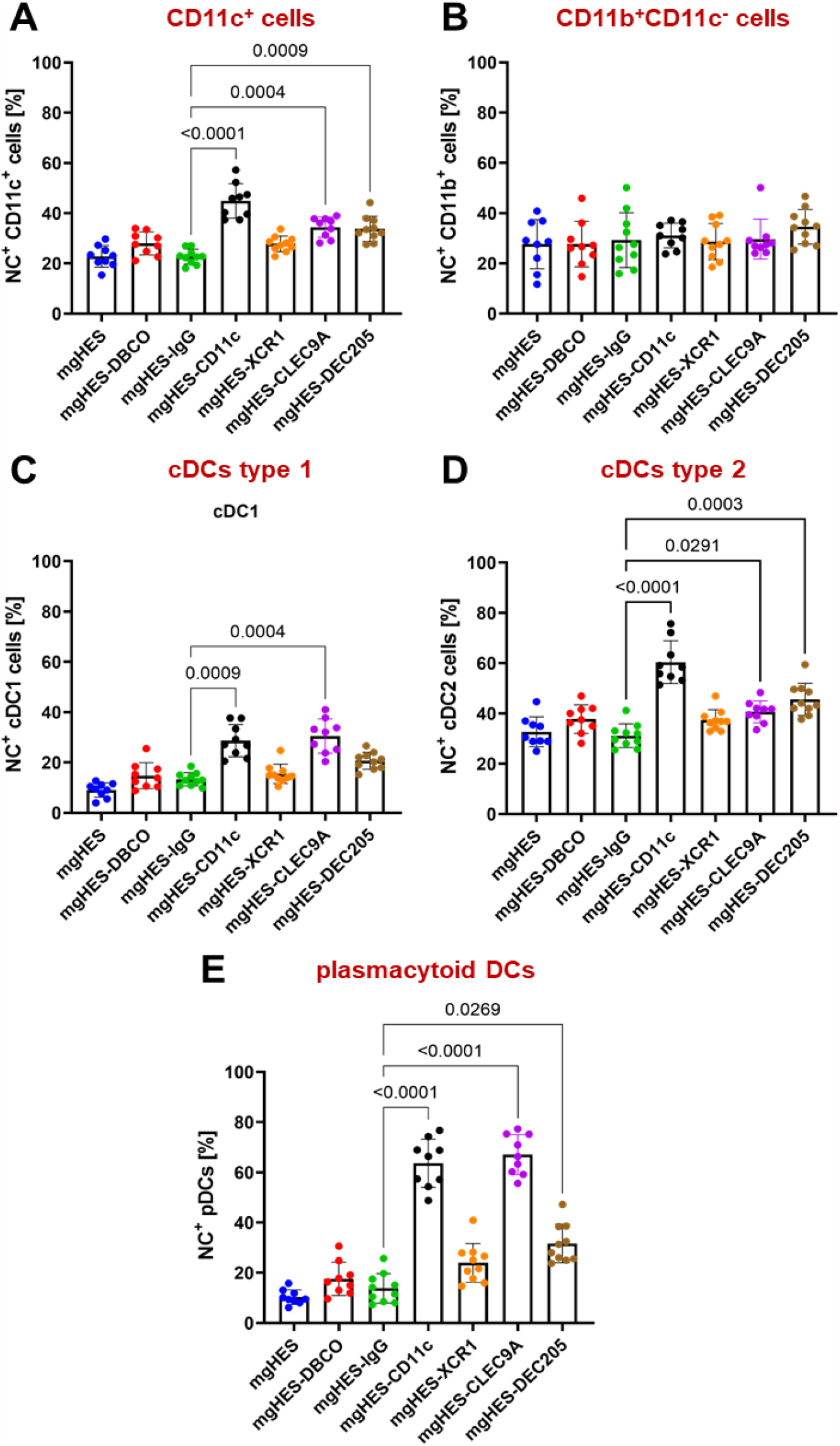
Targeting of dendritic cell subsets *in vivo*. Mice were treated with mgHES nanocarriers functionalized with different DC-targeting antibodies. Nanocarrier uptake by **(A)** CD11c^+^ cells, **(B)** CD11b^+^CD11c^-^ macrophages, **(C)** conventional DCs type 1, **(D)** conventional DCs type 2, and **(E)** plasmacytoid DCs within the spleen was determined using flow cytometry. Data represent mean ± SD. All nanocarrier formulations were compared to the IgG control (mgHES-IgG) and significance was given with *p* < 0.05 using a Kruskall-Wallis test followed by a Dunnett’s multiple comparison test. Individual *p* values are indicated in the graph.

## Conclusion

In this study, dendritic cell-targeting antibodies were successfully conjugated to magnetic nanocarriers in a site-specific and orientated manner. The biomolecular corona that formed onto the nanocarriers after intravenous injection did not prevent binding of the antibodies towards the cell surface receptors on CD11c^+^ dendritic cells *in vitro* and *in vivo*. CD11c and CLEC9A antibodies proved to be excellent candidates for DC targeting, while anti-CLEC9A exhibited a specific targeting towards cDC1 and plasmacytoid DCs, representing the cells of interest. Proteomic analysis revealed major differences in the composition of the protein corona formed during *in vitro* incubation and *in vivo* circulation explaining the great discrepancies of successful *in vitro* and failing *in vivo* studies in terms of antibody-mediated nanocarrier targeting. To overcome this problem, we developed an *ex vivo* approach using whole blood, which allowed us to mimic the protein corona formed *in vivo*. In conclusion, this study presents a detailed protocol for the attachment of antibodies on a nanocarrier surface and demonstrates how to prove the targeting specificity *in vivo*, which can be transferred to a broad range of nanocarriers and antibodies. This novel conjugation technique paves the way for the development of antibody-functionalized nanocarriers for DC-based vaccination approaches in the field of cancer immunotherapy.

## Acknowledgements

This work was partially supported by the German Research Foundation within Sonderforschungsbereich 1066 (SFB 1066).

## Supplementary Information

Material, methods, and additional data is summarized in the supplementary information. A list of all identified proteins is provided in a separate Excel Sheet.

