## Supplementary Information for "Achieving dendritic cell subset-specific targeting *in vivo* by site-directed conjugation of targeting antibodies to nanocarriers"

[1]shared first author

*Corresponding author

### 1. Material and methods

**Nanocarriers.** Hydroxyethyl starch-coated magnetic nanoparticles (BNF-Starch) with amino groups on the surface (named as mgHES-$NH_2$) were obtained by MicroMod. Nanoparticles either contained the fluorophores Dy-750 (for *in vivo* imaging) and Dy-633 (for *in vitro* assays) or redF (for the comparison of different DC-targeting antibodies). Nanoparticles were synthesized via the core-shell method according to literature [1, 2] with a solid content of 5 mg $mL^{-1}$ and 3 nmol $mg^{-1}$ amino groups per mg nanoparticle.

**Enzymatic modification of antibodies.** Antibodies (anti-mouse CD11c (clone N418), anti-mouse CLEC9A (clone 7H11), anti-mouse/rat XCR1 (clone ZET), anti-mouse DEC205 (clone NLDC-145), or IgG isotype controls (Armenian Hamster IgG or Mouse IgG2b, κ) all from Biolegend) were modified with azide groups via the Site Click™ Antibody Azido Modification Kit (Thermo Fisher Scientific) according to the manufacturer's instructions. Briefly, the carbohydrate domain at the Fc region of the concentrated antibody was modified by cleaving the galactose residues using β-galactosidase for 6 h at 37 °C. Afterwards, the azide group was attached by applying the GalT (Y289L) enzyme in combination with the UDP-GalNAz donor in an overnight reaction at 30 °C. The azide-modified antibody was purified via Zeba™ Spin columns, concentrated and the protein concentration was determined.

**Protein quantification.** The total amount of antibody or protein concentrations were quantified by using the Pierce 660 nm protein assay according to the manufacturer's instructions. Bovine serum albumin (BSA) was used as a standard by preparing a dilution series in demineralized water. The optical density at 660 nm of samples and standards was measured in duplicates using an Infinite M1000 plate reader (Tecan).

**Nanocarrier functionalization with antibodies.** mgHES-$NH_2$ nanoparticles (200 mg, 40 mL) were functionalized with either DBCO-$PEG_4$-NHS linker (3.9 mg, 10 x molar excess, 20 mg $mL^{-1}$ stock solution in DMSO; Jena Biosciences) or DBCO-$PEG_{5kDa}$-NHS linker (150 mg, 50x molar excess, 20 mg $mL^{-1}$ stock solution in DMSO; Nanocs Inc., USA). The reaction was incubated overnight at room temperature. The following day, the nanoparticle dispersion was washed three times with 1 mL of PBS by applying a neodymium to separate unbound linker moieties. DBCO-functionalized nanocarriers (mgHES-DBCO, 67 mg) were incubated with azide-modified antibodies (1 mg) at room temperature under shaking overnight. The following day, the dispersion was washed three times with PBS to remove unconjugated antibodies.

**Nanocarrier quantification.** To determine the amount of nanocarrier and nanocarrier-conjugates, a fluorescence calibration of the nanocarrier was conducted. The non-functionalized nanocarriers were applied as a standard and the samples diluted based on the initial mass used. All probes, including the standard, were prepared as duplicates in deionized water. Fluorescence was measured using an Infinite M1000 plate reader (Tecan) with an excitation wavelength of 552 nm or 633 nm and emission of 580 nm or 647 nm for redF-labelled or Dy-633-labelled mgHES nanocarriers, respectively.

**Dynamic light scattering.** The size (diameter in nm) and size distribution (PDI) of all nanocarriers were measured by dynamic light scattering (DLS, Malvern Instruments GmbH, Herrenberg, Germany) at 25 °C at an angle of 90°. The nanocarriers (10 µL, 10 mg $mL^{-1}$) were diluted in PBS and measurements were performed in technical triplicates. Multi-angle light scattering measurements were performed on an ALV spectrometer consisting of a goniometer and an ALV-5004 multiple-tau full-digital correlator (320 channels) at 20 °C and 7 (dynamic) angles ranging from 30° to 150°. A He-Ne Laser (wavelength of 632.8 nm) was used as light source. 1 µl of unfiltered dispersion was diluted with 1 ml PBS which was previously filtered through membrane filters with a pore size of 0.20 µm (GS Millipore).

**Zeta potential.** Measurements were performed with a Zeta Sizer Nano Series (Malvern Instruments GmbH, Herrenberg, Germany) and nanocarriers (10 µL, 10 mg $mL^{-1}$) were diluted in an 1 mM potassium chloride solution (1 mL). Each measurement was performed in technical triplicates.

***In vivo* animal studies.** Wild type C57BL/6 mice and C57BL/6 albino mice (B6N-Tyrc-Brd/BrdCrCrl) were kept in the animal facility of the Translational Animal Research Center at the University Medical Center of the Johannes Gutenberg University Mainz under pathogen-free conditions on a standard diet. All procedures were approved by the regional animal care committee and performed according to the EU Directive 2010/63/EU. Nanocarriers (1 mg in 500 µL PBS or 500 µg in 200 µl PBS (for comparing different antibody-functionalized nanocarriers) were administered intravenously via tail vein injection. As a control, animals were treated with PBS. After 1 min, 10 min, 60 min, or 120 min blood was isolated via cardiac puncture. Animals were sacrificed and all organs (lung, spleen, liver, and kidney) were prepared and imaged via small animal fluorescence imaging (IVIS® SpectrumCT, Perkin Elmer). For the comparison of different DC-targeting antibodies conjugated onto the nanocarrier surfaces, mice were sacrificed 24 h after injection via cervical dislocation and spleens were dissected and dissociated for subsequent flow cytometric analyses.

**Isolation of spleen cells.** Spleen cells were prepared as previously described[3]. Briefly, the spleens of C57BL/6 mice were isolated and mechanically disrupted by grinding through a 40 µm cell strainer. Subsequently, erythrocytes within the cell suspension were lysed using a hypotonic buffer (155 mM $NH_4Cl$, 10 nM KHCO3, 100 µM EDTA-disodium, pH 7.4) for 30 s and subsequently washed using Iscove's Modified Dulbecco's Medium (IMDM) supplemented with 5% fetal bovine serum (FBS) and 1% Penicillin (100 U $mL^{-1}$) and Streptomycin (100 mg $mL^{-1}$ (P/S). Spleen cells were stained with antibodies to discriminate the different cell populations and the amount of nanocarrier positive cells was determined by flow cytometry.

**Spleen cell staining.** First, cells were washed (1% FBS in PBS) and Fc receptors were blocked with anti-CD16/CD32 (clone 2.4G2) for 10 min. To differentiate the cell types via flow cytometry, spleen cells were incubated with fluorophore-labelled cell type-specific antibodies for 30 min: anti-CD11c (clone N418), anti-CD11b (clone M1/70), anti-CD172a (clone P84), anti-CD8α (clone 53-6.7), anti-I-A/I-E (MHCII, clone M5/114.15.2), anti-F4/80 (clone BM8), anti-CD4 (clone GK1.4), anti-Siglec-H (clone 551), anti-CD19 (clone 6D5), anti-CD3ε (clone 145-2C11), anti-CD14 (clone Sa14-2), anti-NK1.1 (clone PK136), anti-Ly6G (clone 1A8).Flow cytometric analysis was performed with the Attune NxT (Thermo Fisher Scientific) and analyzed with FlowJo software v10.7.1.

**Isolation of liver cells.** Non-parenchymal liver cells of C57BL/6 mice were isolated by an organ perfusion method as previously described [4]. Therefore, mice were anesthetized using ketamine/xylazine and the abdominal cavity was opened. The liver was flushed through the portal vein with 20 ml of $Ca^{2+}$- and $Mg^{2+}$-free HBSS (Hank's Balanced Salt Solution, Thermo Fisher Scientific) containing 5% FBS, collagenase A (100 U $L^{-1}$; Sigma-Aldrich) and DNase I (10 µg $mL^{-1}$; Sigma-Aldrich). The liver was subsequently cut into pieces, incubated in the perfusion solution for 15 min at 37 °C in a 50 ml tube, and mechanically disrupted with a 70 µm nylon cell strainer. The enzymatic reaction was stopped by adding a cell culture medium (Dulbecco's Modified Eagle Medium (DMEM)/F-12, 5% FBS, 1% P/S). Hepatocytes were pelleted and removed via centrifugation (30 g for 15 min) and the remaining cells in the supernatant were purified by 30% Histodenz-HBSS gradient centrifugation according as previously described [5]. Non-parenchymal liver cells were obtained, stained with antibodies, and the frequency of nanocarrier positive cells was determined by flow cytometry.

**Liver cell staining.** First, cells were washed (1% FBS in PBS) and Fc receptors were blocked with anti-CD16/CD32 (clone 2.4G2) for 10 min. To discriminate the cell types, liver cells were incubated with cell type-specific antibodies for 30 min: anti-CD45 (clone 30-F11), anti-CD31 (clone 390), anti-CD11c (clone N418) and anti-F4/80 (clone BM8)

***In vivo* protein corona.** Nanocarriers were recovered from the blood stream 1 min, 10 min,

60 min, or 120 min after intravenous injection via cardiac puncture followed by magnetic separation. The resulting nanocarrier pellet was washed with PBS (three times) to remove loosely bound and unbound proteins. Afterwards, the firmly attached proteins were desorbed from the surface using 2% SDS (with 62.5 mM Tris*HCl) and heated up to 95 °C for 5 min. The nanocarrier pellet was magnetically separated and the resulting protein supernatant was analyzed by Pierce Assay, SDS-PAGE, and liquid chromatography-mass spectrometry (LC-MS).

***Ex vivo* protein corona.** Blood was isolated from C57BL/6 mice via cardiac puncture (~500 µL – 1 mL) and supplemented with heparin (2 µL, 5,000 units). Nanocarriers (1 mg) were incubated with 2 mL of whole blood for 1 min and the *ex vivo* protein corona was purified as described above (*in vivo* protein corona).

***In vitro* protein corona.** Mouse serum and plasma, derived from anti-coagulation with EDTA, heparin, or citrate, was prepared from C57BL/6 mice blood. Nanocarriers (1 mg) were incubated with serum or plasma (1 mL) for 1 min at 37 °C. The *in vitro* protein corona around the nanocarriers was purified as described above (*in vivo* protein corona).

**SDS-PAGE.** Corona proteins (1 - 2 µg) were mixed with NuPAGE™ LDS sample buffer and NuPAGE™ sample reducing agent, incubated at 95 °C for 5 min, and further applied to a NuPAGE™ 10% Bis-Tris protein gel (Thermo Fisher Scientific). The gel was run in NuPAGE™ MES SDS running buffer at 120 V for 1 h with dithiothreitol (DTT) and SeeBlue Plus2 Pre-Stained Standard was used as a molecular marker. Proteins were visualized using the SilverQuest™ Silver Staining kit (Thermo Fisher Scientific) according to the manufacturer´s instruction.

**In solutions digestion.** Proteins were digested according to previous reports [6, 7]. First, SDS was removed via Pierce detergent removal columns (Thermo Fisher Scientific). Further, proteins were precipitated overnight using the ProteoExtract® Protein Precipitation Kit (Millipore) according to the manufacturer’s instruction. The protein pellet was re-suspended in an ammonium bicarbonate (50 mM) buffer supplemented with RapiGest SF (Waters Cooperation). Subsequently, proteins were reduced (DTT, 5 mM, Sigma-Aldrich) for 45 min at 56 °C, alkylated with (iodoacetoamide, 15 mM, Sigma-Aldrich) for 60 min at room temperature and finally trypsin was added. A protein:trypsin ratio of 50:1 was used and the digestion was carried out for 14 h to 18 h at 37 °C. The reaction was quenched with 2 µL hydrochloric acid (Sigma-Aldrich).

**Liquid chromatography coupled to mass spectrometry (LC-MS).** The reference peptide standard Hi3 E.coli (Waters Cooperation) was spiked into all peptide samples at a concentration of 50 fmol $\mu L^{-1}$ for absolute protein quantification [8]. Measurements were performed with a nanoACQUITY UPLC coupled to a Synapt G2-Si mass spectrometer (Waters Cooperation). Ionization was carried out with a NanoLockSpray source in positive ion mode and the Synapt G2-Si was operated in resolution mode performing data-independent acquisition ($MS^E$) experiments. For data analysis MassLynx 4.1 and Progenesis QI (2.0) was used. A murine database was downloaded from Uniprot. Proteins were quantified based on the TOP3/Hi3 [9] providing the absolute amount of each protein in fmol. For further analysis, the relative amount in % based on all identified proteins was calculated.

**Cell culture.** Murine DC2.4 dendritic cells (Merck) were cultured in IMDM supplemented with 5% FBS, 1% P/S, and 50 µM 2-Mercaptoethanol at 37 °C and 5% $CO_2$. Cells were regularly split (2-3 times per week) when reaching a confluence of about than 80%.

***In vitro* DC2.4 cell uptake.** Cells (100.000 per well) were seeded in a 24-well plate with a regular cell culture medium containing FBS and incubated overnight. The next day, cells were washed with PBS and FBS-free medium was added (1 mL per well). Cells were incubated for 2 h. Nanocarriers left untreated, pre-treated with mouse serum/plasma (ratio see *in vitro* protein corona) or isolated after *in vivo* intravenous injection were added to cells at a concentration of 37.5 µg $mL^{-1}$ in cell culture medium without FBS for 2 h. Cells were washed with 1 mL PBS and detached with 2 mM EDTA in PBS. After centrifugation (300 x g, 5 min), the resulting cell pellet was resuspended in 1 mL of PBS + 2% FBS and analyzed by flow cytometry using the

Attune™ NxT (Thermo Fisher Scientific). Cells were selected on a FSC/SSC scatter plot, excluding cell debris populations and the median fluorescence intensity (MFI) or as the percentage of gated events/cells. The red fluorescence of the nanocarriers was detected by the RL1 channel with an excitation laser of 638 nm and a 670/14 nm band pass emission filter.

***Ex vivo* nanoparticle-blood interaction.** Blood was isolated from C57BL/6 albino mice via cardiac puncture (~500 µL - 1 mL) and supplemented with heparin (2 µl, 5,000 units). Nanocarriers (5 µL, 50 µg) were incubated with whole blood (1 µL) for 1 min. The NP-blood mixture was diluted with 1 mL of PBS and analyzed by flow cytometry using the NxT No-Wash No-Lyse Filter Kit (Thermo Fisher Scientific). Therefore, the Attune NxT flow cytometer was equipped with an additional side scatter channel off of the violet laser, which allowed the immunophenotyping of whole blood without the lysis of red blood cells and antibody staining.

### 2. Protein corona analysis

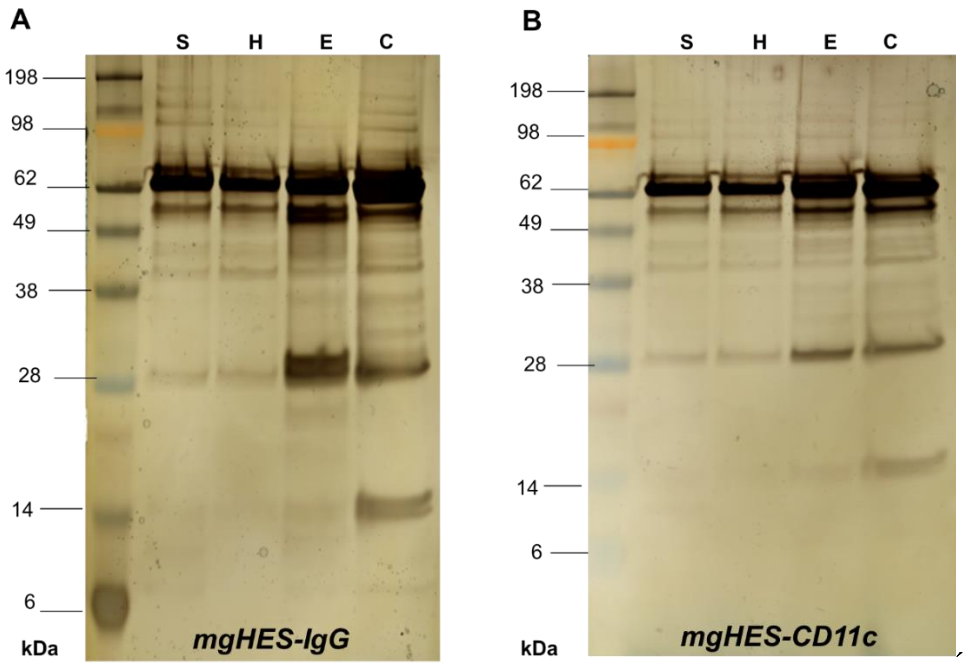

**Figure S1.** SDS-PAGE of the protein corona of antibody-functionalized nanocarriers incubated under *in vitro* conditions for 1 min (S = Serum, H = Heparin plasma, E = EDTA plasma, C = Citrate plasma). **A)** mgHES-IgG nanocarriers vs. **B)** mgHES-CD11c nanocarriers

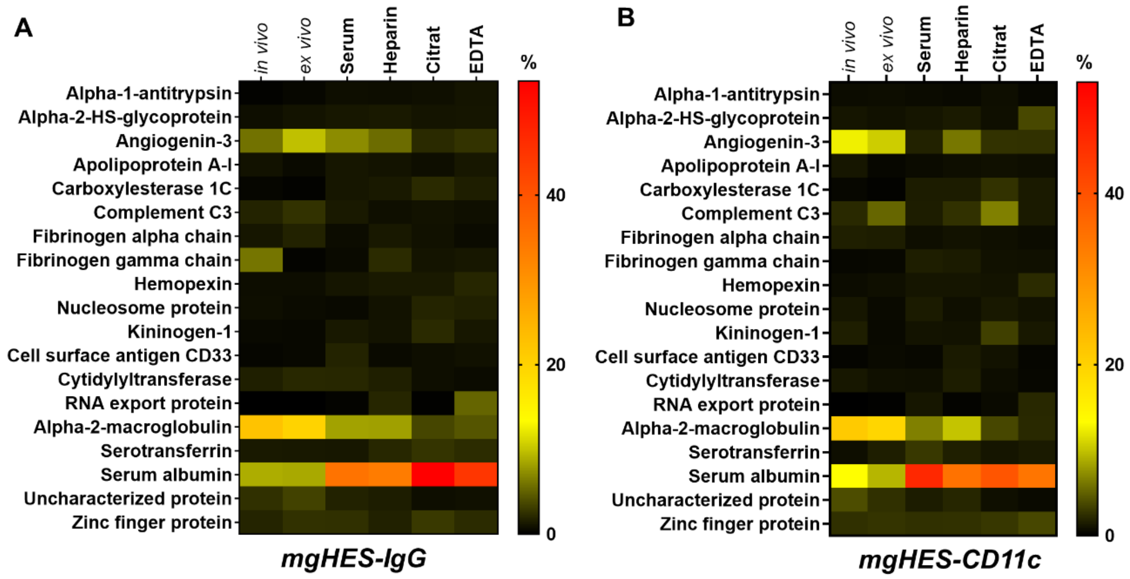

**Figure S2.** LC-MS analysis of the protein corona. The relative amount of each protein in % is calculated based on the total amount of all identified proteins determined in fmol (n = 3, for all *in vivo* samples). **A)** mgHES-IgG nanocarriers vs. **B)** mgHES-CD11c nanocarriers.

### 3. *In vivo* cell targeting

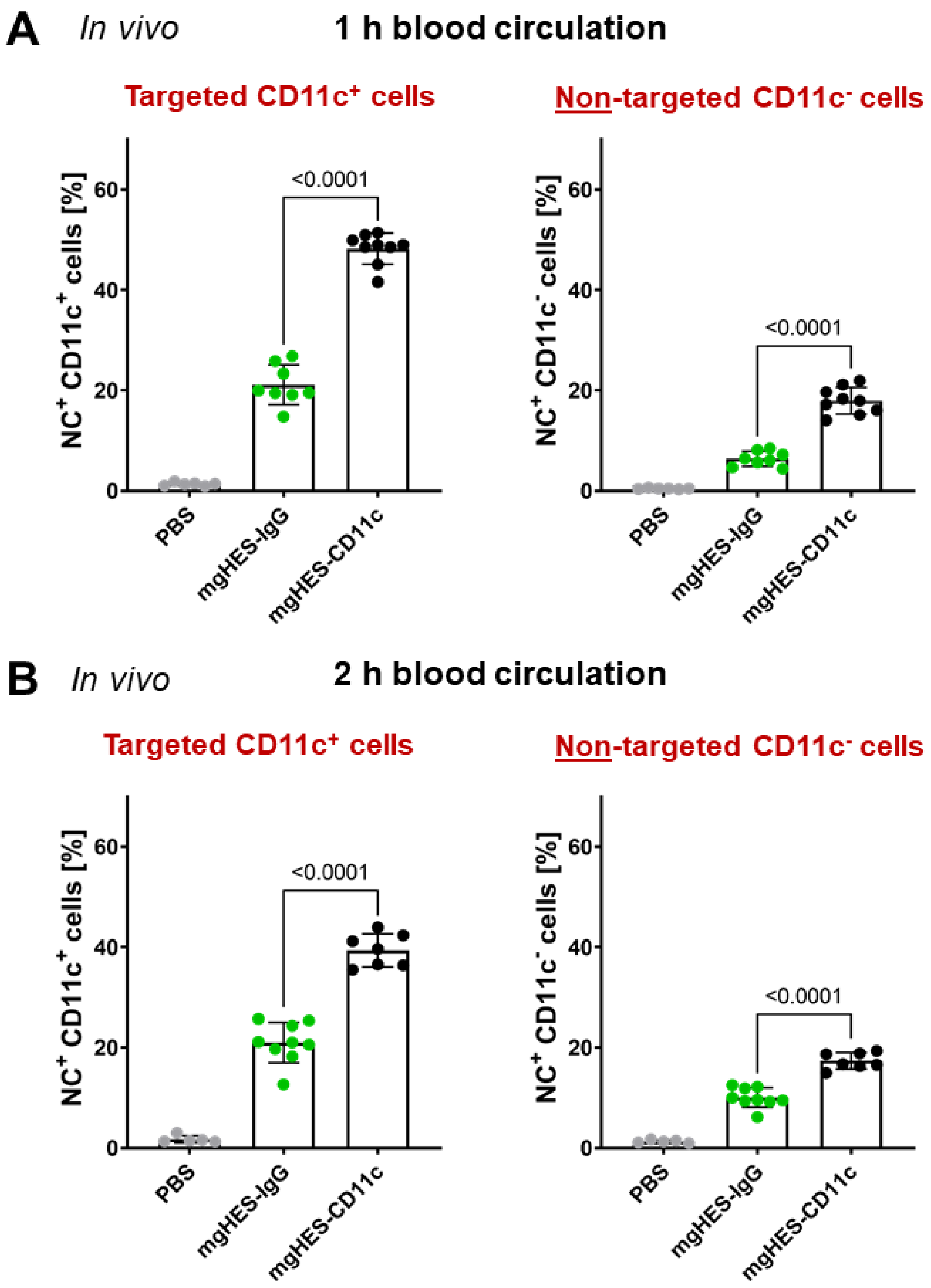

**Figure S3**. Targeting CD11c$^+$ spleen cells *in vivo*. Mice were treated with antibody-conjugated nanocarriers for 1 h **(A)** or 2 h **(B)**, subsequently sacrificed and the spleen was dissected. Binding of antibody-conjugated nanocarriers towards CD11c$^+$ and CD11c$^-$ cells was analyzed by flow cytometry. Data represent mean ± SD. mgHES-CD11c nanocarriers were compared to the IgG control (mgHES-IgG) and significance was given with $p < 0.05$ using a

Welch's *t* test. Individual *p* values are indicated in the graph.

### 4. Dynamic light scattering and zeta potential measurement

Table S1: Physicochemical characterization of antibody-functionalized mgHES nanocarrier.

| Sample name | Average diameter in PBS[a] | PDI in PBS[a] | ξ-potential [mV] | ξ-deviation [mV] |
|---|---|---|---|---|
| mgHES-$NH_2$ | 290 nm | 0.161 | -2.21 ± 0.88 | 3.23 ± 0.17 |
| mgHES-$PEG_{5kDa}$-DBCO | 396 nm | 0.233 | -4.22 ± 0.40 | 3.75 ± 0.34 |
| mgHES-$PEG_{5kDa}$-IgG | 238 nm | 0.129 | -4.63 ± 0.25 | 3.28 ± 0.26 |
| mgHES-$PEG_{5kDa}$-CD11c | 208 nm | 0.076 | -3.84 ± 0.10 | 3.46 ± 0.42 |
| mgHES-$PEG_{5kDa}$-XCR1 | 288 nm | 0.145 | -4.79 ± 0.22 | 4.43 ± 0.50 |
| mgHES-$PEG_{5kDa}$-CLEC9A | 308 nm | 0.186 | -4.85 ± 0.37 | 4.46 ± 0.81 |
| mgHES-$PEG_{5kDa}$-DEC205 | 220 nm | 0.194 | -6.82 ± 0.58 | 5.12 ± 1.31 |

[a] Determined by multi angle DLS (at scattering angles of 30° to 150°)
